# The pERKs of Temporal Order Memory in mice

**DOI:** 10.64898/2026.07.02.736134

**Authors:** Santiago D’hers, Santiago Ojea Ramos, Agustina Denise Robles, Mariana Feld

## Abstract

Temporal Order Memory (TOM), the ability to discriminate between events according to when they occurred, is a key component of episodic-like memory. Understanding the molecular mechanisms that support temporal memory requires behavioral approaches capable of capturing the continuous dynamics of natural exploration. Despite extensive evidence implicating the prefrontal cortex (PFC) in temporal memory, the intracellular signaling mechanisms supporting temporal order discrimination remain poorly understood. Here, we combined high-resolution automated behavioral phenotyping with molecular analyses to characterize the behavioral and signaling dynamics underlying TOM in m|ice.

Mice were trained in a spontaneous object-recognition TOM task and tested after short-term (3 h) or long-term (24 h) retention intervals. Exploration was quantified using an artificial intelligence–based behavioral analysis pipeline that enables continuous and unbiased assessment of object exploration. Phosphorylation of extracellular signal-regulated kinase 2 (ERK2) and expression of the ERK phosphatase MKP3/DUSP6 were analyzed in the PFC and hippocampus (HIP) following habituation, a single training session, or two sequential training sessions. Additionally, a Temporal Novel Object Recognition (TeNOR) protocol was used to evaluate the integrity of memory traces.

Mice displayed robust TOM performance across sexes and retention intervals. Molecular analyses revealed no significant changes in hippocampal ERK signaling, whereas the cytosolic fraction of the PFC exhibited dynamic, experience-dependent modulation of ERK2 phosphorylation. A single 15-minute training session induced a transient increase in ERK2 activation, while a second session 45 minutes later actively suppressed this peak. This rapid molecular reset was accompanied by increased MKP3 expression, suggesting the targeted recruitment of an active regulatory feedback mechanism. Continuous behavioral tracking further revealed temporal features of memory expression that were not captured by conventional summary measures; it identified an early, rapid decay of discrimination for older object memories in the TeNOR task, suggesting that TOM performance relies on resolving competitive retrieval between co-existing memory traces.

Together, these findings identify dynamic ERK2-MKP3 signaling in the PFC as the molecular substrate upon which temporal discrimination can take place, and demonstrate how high-resolution phenotyping in naturalistic behavioral paradigms can reveal mechanistic links between intracellular signaling and the temporal organization of experience.

## 1. Introduction

Understanding how the brain organizes experiences across time is a central challenge in memory research. Episodic memory requires not only recognizing past events, but also encoding their temporal relationships (Eichenbaum, 2017; Tulving, 2002). In rodents, Temporal Order Memory (TOM) paradigms have been widely used to study episodic-like temporal discrimination by evaluating the ability to distinguish between familiar objects encountered at different moments in time (Barker et al., 2007; Dere et al., 2018; Mitchell, 1998). Successful TOM performance is typically inferred from the preferential exploration of an object experienced earlier relative to one encountered more recently (Barker & Warburton, 2011). Spontaneous object-recognition paradigms, such as TOM, provide a naturalistic and minimally stressful framework for studying memory, as they rely on the animal’s intrinsic exploratory drive and novelty preference rather than appetitive or aversive reinforcements (Antunes & Biala, 2012; Ennaceur & Delacour, 1988).

While ethologically relevant, spontaneous object-recognition paradigms pose important analytical challenges because exploratory behavior is inherently dynamic and continuous. Recent advances in automated behavioral quantification, pose-estimation tools, and machine-learning approaches have transformed the study of animal behavior by enabling automated, high-resolution quantification of spontaneous exploration (Datta et al., 2019; Mathis et al., 2018). These methods move beyond traditional manual analysis, which is often prone to subjective bias and limited reproducibility, and capture the continuous dynamics of behavior with greater precision and scalability.

In this context, we recently developed RAINSTORM (Real and Artificial Intelligence for Neuroscience — Simple Tracker for Object Recognition Memory; D’hers et al., 2025), an automated analysis pipeline that combines pose-estimation data, geometric analysis, and artificial intelligence–based behavioral classification to quantify exploratory behavior in rodents. By learning from expert-defined scoring criteria, RAINSTORM minimizes evaluator bias while providing rapid and accurate assessment of object exploration and recognition memory performance. In the present study, we used RAINSTORM to assess TOM in mice, enabling high-resolution analysis of temporal memory expression and exploratory dynamics.

The hippocampus (HIP) and prefrontal cortex (PFC) have both been implicated in temporal memory and object recognition processes (Barker & Warburton, 2011). However, their functional contributions are distinct: while the HIP is crucial for the integrated ‘what-where-when’ components of episodic memory (Clayton et al., 2001), the PFC specifically supports the organization of sequential information and the representation of temporal relationships between events (DeVito & Eichenbaum, 2011). Damage to the PFC often results in a ‘disordered’ memory, characterized by the ability to recognize an object (the ‘what’) but a fundamental failure to recollect its specific temporal (the ‘when’) or spatial (the ‘where’) context (Barker et al., 2007; Janowsky et al., 1989). Despite this circuit-level understanding, the intracellular signaling mechanisms supporting these computations remain poorly understood.

One candidate mechanism for TOM processing is the extracellular signal-regulated kinase (ERK1/2) pathway, a central regulator of synaptic plasticity, learning, and long-term memory formation (Sweatt, 2004; Thomas & Huganir, 2004). ERK signaling is dynamically modulated during learning and novelty exposure across multiple species and memory paradigms (Adams & Sweatt, 2002; Antoine et al., 2014). Importantly, previous work in *Drosophila* proposed that repeated learning events can reset ERK activation states, potentially preventing molecular overlap between sequential memories (Cattaneo et al., 2020). Similarly, studies in the crab *Neohelice granulata* demonstrated that ERK signaling is critically involved in the formation of a long-term memory induced by a two-trial training protocol with a 45-min interval, further supporting the idea that tightly regulated ERK dynamics may contribute to memory formation under sequential learning conditions (Ojea Ramos et al., 2021). Whether similar mechanisms contribute to TOM processing in mammals remains unknown.

The duration and magnitude of ERK activation are tightly regulated by phosphatases such as MKP3/DUSP6, a cytosolic dual-specificity phosphatase that negatively regulates ERK1/2 signaling (Caunt & Keyse, 2013; Farooq & Zhou, 2004). Although ERK signaling has been extensively associated with synaptic plasticity and memory processes, the contribution of its regulatory phosphatases (MKP3 in particular) remains largely unexplored. Importantly, previous work from our group has consistently highlighted the relevance of cytosolic ERK dynamics during long-term memory consolidation and reconsolidation across different experimental models (Feld et al., 2005, 2008; Krawczyk et al., 2015, 2016; Ojea Ramos et al., 2026). These findings suggest that precise regulation of cytosolic ERK activity is critical for memory processing. Dynamic interactions between ERK phosphorylation and MKP3-mediated dephosphorylation could therefore provide an active regulatory mechanism to limit molecular overlap and compartmentalize intracellular signaling during sequential learning. Nevertheless, the role of MKP3 in TOM has not yet been investigated.

Another unresolved question in TOM research is whether the preference for the “older” object reflects genuine temporal discrimination or an emergent property of competitive retrieval and retroactive interference (Autore et al., 2023). Temporal Novel Object Recognition (TeNOR) paradigms provide an opportunity to dissociate these possibilities by independently probing the recognition of old versus recent familiar objects.

In the present study, we combined automated behavioral quantification with spontaneous object-recognition paradigms to characterize the behavioral and molecular dynamics underlying TOM in mice. Specifically, we analyzed short- and long-term TOM performance, characterized ERK2 phosphorylation dynamics and MKP3 expression in the PFC and HIP during memory acquisition, and used the TeNOR paradigm to investigate the relative stability of old versus recent memory traces. Our results identify a dynamic ERK2 and MKP3 signaling profile in the PFC associated with sequential learning and suggest a potential molecular mechanism for temporal discrimination between consecutive experiences.

## 2. Materials and methods

### 2.1. Animals

Male and female C57BL/6 elite mice (obtained from the *Academia Nacional de Medicina*, Buenos Aires, Argentina) were used for all experiments. A total of 107 mice, starting at 8 to 9 weeks of age, were employed across different experimental cohorts, as detailed in Table 1. To mitigate potential carryover effects from repeated testing, the assignment of objects and their spatial locations were fully counterbalanced across all training and testing sessions. Furthermore, subjects were exposed to a unique set of novel objects for each session, ensuring no object was repeated for any experimental subject.

**Table 1:** Summary of experimental cohorts included in this study. Distribution of mice assigned to behavioral and molecular experiments, detailing total sample sizes (n), sex distribution, and animal age at the beginning of the experimental protocols.

| Experiment | Cohort | Age | Mice (n) | Females | Males |
| --- | --- | --- | --- | --- | --- |
| Short-term TOM | A | 8 weeks | 23 | 11 | 12 |
| Short-term TOM |  | 12 weeks |  |  |  |
| Long-term TOM |  | 14 weeks |  |  |  |
| Long-term TOM | B | 8 weeks | 20 | 12 | 8 |
| TeNOR |  | 10 weeks |  |  |  |
| Molecular characterization | C | 8 weeks | 64 | 25 | 39 |
| **Total** |  |  | **107** | **48** | **59** |

Mice were group-housed in plastic cages (2 to 4 individuals per cage) under standard laboratory conditions: controlled temperature (24° ± 2°), a 12-hour light-dark cycle (lights on at 07:00), and *ad libitum* access to food and water. All animal handling and experimental procedures were performed in strict accordance with the NIH Guide for the Care and Use of Laboratory Animals and approved by the Institutional Animal Care and Use Committee (CICUAL – FCEN, UBA, protocols N° 165/2021 and 165b/2025).

### 2.2. Object Recognition Protocols

Behavioral tasks were conducted in a white polyurethane open-field arena (20 x 30 cm). The arena floor was removable and equipped with four hidden magnets situated 5 cm from each corner, designed to securely anchor the objects and prevent displacement during exploration. The objects included blue LEGO bricks, glass jars, cylindrical plastic objects, and ping-pong balls. All objects were previously validated in our laboratory to ensure the absence of any innate bias or intrinsic preference among mice.

#### 2.2.1. Temporal Order Memory (TOM)

The TOM protocol was composed of the following phases:

- Habituation (HAB): Three sessions on consecutive days (5, 10, and 15 min), during which each mouse was placed in the empty arena and allowed to freely explore it.
- First training session (TR1): On the fourth day, the mouse was placed in the arena containing two identical objects and was allowed to explore for 15 min.
- Second training session (TR2): After a 45-min interval, the mouse was put into the arena again to explore a new pair of identical objects for 15 min.
- Testing session (TS): 3 h (short-term) or 24 h (long-term) after the end of the second training, the animal entered the arena to explore one object from TR1 (old familiar object) and one from TR2 (recent familiar object) for 5 min. TOM is defined by the preference for exploring the old familiar object, known from TR1, over the recent familiar one (presented at TR2).

#### 2.2.2. Temporal Novel Object Recognition (TeNOR)

The arena, objects, and protocol are identical to the TOM protocol, except for the TS session.

- Testing session (TS): 24 h after the end of the TR2 session, mice were divided into two groups. The first group (Prior_TR_) entered the arena to explore one object from TR1 and a novel object, while the second group (Latter_TR_) explored one object from TR2 and a novel object. In both cases, recognition memory is defined by the preference for exploring the novel object over the familiar (old from TR1, or recent from TR2) one.

In both protocols, discrimination performance was expressed as a Discrimination Index (DI), calculated as the proportion of exploration time directed toward the target object (i.e., the older object in TOM and the novel object in TeNOR) relative to the total object exploration time during the first 4 minutes of TS.

### 2.3. Automated Quantification of Exploratory Behavior

Exploratory behavior was recorded using a Genius ECAM-8000 webcam interfaced with Bonsai software (version 2.8.5; Lopes et al., 2015). Body-part tracking was performed using DeepLabCut (version 3.0.0; Mathis et al., 2018; Nath et al., 2019). Specifically, we manually labeled 300–350 frames extracted from 30 representative videos, allocating 95% of the frames for training and the remaining 5% for testing. We trained a ResNet-50-based neural network for 800 epochs (test error: 3.02 px, train error: 1.39 px; image resolution: 720 x 640 px). The trained network was subsequently applied to extract positional data from all experimental videos. The resulting coordinates (nose, head, neck, ears, body center, and tail base) were processed using the RAINSTORM pipeline (D’hers et al., 2025), which leverages artificial neural networks to automate the quantification of exploratory behavior.

### 2.4. Tissue Preparation and Western Blotting

To evaluate molecular dynamics, mice were euthanized at specific time points (0, 1, and 2 h) following either HAB, TR1, or TR2 (the full two-session training protocol). Following euthanasia by cervical dislocation, brain tissue from the HIP and PFC regions was rapidly dissected, and extracts enriched in nuclear or cytosolic proteins were prepared as described in Salles et al. (2015). Briefly, HIP or PFC tissue was homogenized in 250 μL or 150 μL, respectively, of buffer A (Table 2), with 8 strokes using a glass-glass Dounce homogenizer with a type B (tight) pestle. The homogenate was centrifuged for 15 min at 1000 g, and the supernatant (cytosolic protein-enriched extract) was aliquoted and stored at −20 °C until use. HIP or PFC pellets were resuspended in 50 μL or 20 μL, respectively, of buffer B (Table 2) and incubated on ice for 20 min. They were then centrifuged for 15 min at 12,000 g. The supernatant (nuclear protein-enriched extract) was aliquoted and stored at −20 °C until use. The entire extraction protocol was performed at 4 °C.

**Table 2:** Solutions used throughout this study. *, Phosphatase and protease inhibitors (added freshly the day of use): 1 µg/ml pepstatin A, 10 µg/ml leupeptin, 0.5 mM PMSF, 10 µg/ml aprotinin, 1 mM sodium orthovanadate, 50 mM NaF.

| Buffer | Composition |
| --- | --- |
| A | 10 mM HEPES pH 7.9, 10 mM KCl, 1.5 mM MgCl<sub>2</sub>, 1 mM DTT, *. |
| B | 20 mM HEPES pH 7.9, 800 mM KCl, 1.5 mM MgCl<sub>2</sub>, 0.4 mM EDTA, 0.5 mM DTT, 50 % glycerol, *. |
| Resolving gel 12 % | 4 mL deionized water, 3.3 mL of 29:1 acrylamide:bis-acrylamide solution, 2.5 mL of 1.5 M Tris-HCl (pH 8.8), 100 μL of 10 % SDS, and (right before pouring the solution) 100 μL of 10 % APS, 10 μL TEMED. |
| Stacking gel | 6 mL deionized water, 1.3 mL of 29:1 acrylamide:bis-acrylamide solution, 2.5 mL of 0.5 M Tris-HCl (pH 6.8), 100 μL of 10 % SDS, and (right before pouring the solution) 100 μL of 10 % APS, 10 μL TEMED. |
| Running | 25 mM Tris-Cl, 250 mM glycine (pH 8.3), 0.1 % SDS. |
| Transfer | 25 mM Tris-Cl, 250 mM glycine (pH 8.3), 10 % Methanol. |
| Loading (4x) | 250 mM Tris-HCl (pH 6.8), 8 % SDS, 8 % beta-mercaptoethanol, 0.002 % bromophenol blue, 40 % glycerol. |

Protein concentration was quantified using the Bradford method (Bradford, 1976), with bovine serum albumin (BSA) standards of known mass serving as references. Absorbance was measured at 595 nm in triplicate, and the concentration of each sample was calculated using the linear portion of the BSA calibration curve. Samples (20–30 µg of total protein) were denatured in loading buffer (see Table 2) by heating for 5 min at 95°C, and were resolved by SDS-denaturing polyacrylamide gel electrophoresis (SDS-PAGE) for 20 min at 80 V, and 1–1.5 h at 120 V. Proteins were then electrotransferred under 350 mA for 1 h to polyvinylidene fluoride (PVDF) membranes.

Membranes were left to dry out and reactivated in methanol for a few seconds before blocking for 1 h with 4 % fat-free powdered milk in TTBS (TBS - 0.1 % Tween-20). After blocking, the membranes were incubated with primary antibodies overnight at 4 °C, and then with secondary antibodies at room temperature for 1 h. The following primary antibodies were used: phospho-ERK1/2 (1:1000, Santa Cruz Biotechnology cat. sc-7383, mouse monoclonal), total ERK1/2 (1:1000, Cell Signaling Technology cat. #9102, rabbit polyclonal), and total MKP3 (1:1000, Santa Cruz Biotechnology cat. sc-377070, mouse monoclonal). Total MKP3 was detected by chemiluminescence (Anti-Mouse IgG, Cell Signaling cat. #7076, 1:5.000) using Luminol reagent (ECL, BioRad, cat. #170-5060) and an Amersham Imager 600. Total and phospho-ERK1/2 were detected using IRDye subclass-specific secondary antibodies (donkey anti-mouse, 1:10.000, Li-Cor cat. #926-32212; donkey anti-rabbit, 1:10.000, Li-Cor cat. #926-68073) on an Odyssey imaging system. The images obtained were analyzed using ImageJ 1.53j (Schneider et al., 2012) and GelAnalyzer (version 26.1; Istvan Lazar, 2026), and the signal intensity was determined for the specific bands of either phosphorylated or total ERK1 and ERK2, and total MKP3.

### 2.5. Statistical Analysis

Data processing and statistical analyses were conducted using R (version 4.6.0; R Core Team, 2026) and RStudio (version 2026.05.0-218; Posit team, 2026). To robustly model our experimental data, which included bounded proportions and non-normally distributed continuous variables, we employed Generalized Linear Mixed-Effects Models (GLMMs).

Behavioral discrimination data (DI) from the TOM and TeNOR tasks were modeled using a Beta regression family with a logit link function via the *glmmTMB* package (version 1.1.14; Brooks et al., 2017). This approach is mathematically appropriate for variables strictly bounded between 0 and 1; notably, all calculated DI values fell within this range, as every mouse explored each object at least once per session. The primary model formula incorporated fixed effects for group, sex, and their interaction. To account for repeated measures and control for environmental factors, a parsimonious random effects structure was applied using random intercepts for individual subjects nested within their homecage identity. Finally, the performance package (version 0.12.2; Lüdecke et al., 2021) was used to calculate the intraclass correlation coefficient (ICC) to assess the variability associated with these random intercepts. If the random intercept for “Cage” resulted in a variance of zero (a singular fit), the cage effect was considered negligible.

Molecular data exhibited a continuous, right-skewed positive distribution and were therefore analyzed utilizing a Gamma regression family with a log link function. The model evaluated the interacting fixed effects of training condition and wait time. To account for non-experimental variance and the nested structure of the data, random intercepts for sex and home-cage identity were included.

Following model specification, assumptions for all GLMMs (including residual uniformity, absence of overdispersion, and outlier detection) were evaluated and confirmed using simulated residuals via the *DHARMa* package (version 0.4.7; Hartig, 2026). Main effects and interactions were assessed using Analysis of Variance (ANOVA). Type II ANOVA was explicitly utilized for the molecular data to account for the asymmetric experimental design (specifically, the absence of the HAB_1h_ and TR2_0h_ groups), whereas Type III tests were applied to balanced models. Finally, post-hoc pairwise comparisons and targeted contrasts—against the chance level (DI = 0.5) for behavior, and against the global baseline (HAB_0h_) for molecular assays—were conducted using estimated marginal means (*emmeans* package, version 1.10.3; Lenth & Piaskowski, 2026). Statistical significance was defined at α = 0.05.

## 3. Results

### 3.1. Behavioral characterization of TOM formation

First, we characterized the formation of Temporal Order Memory (TOM) using a behavioral protocol consisting of two training sessions (TR1 and TR2) separated by a 45-min interval, during which mice were exposed to different pairs of identical objects. The temporal order memory was then tested (TS) at short-term (3 h) or long-term (24 h) intervals after TR2. During TS, mice were presented with one object from each training session (Figure 1A). Given that both objects were familiar, successful TOM was inferred when mice preferentially explored the object presented earlier in time (old familiar object) over the one presented more recently.

**Figure 1:**
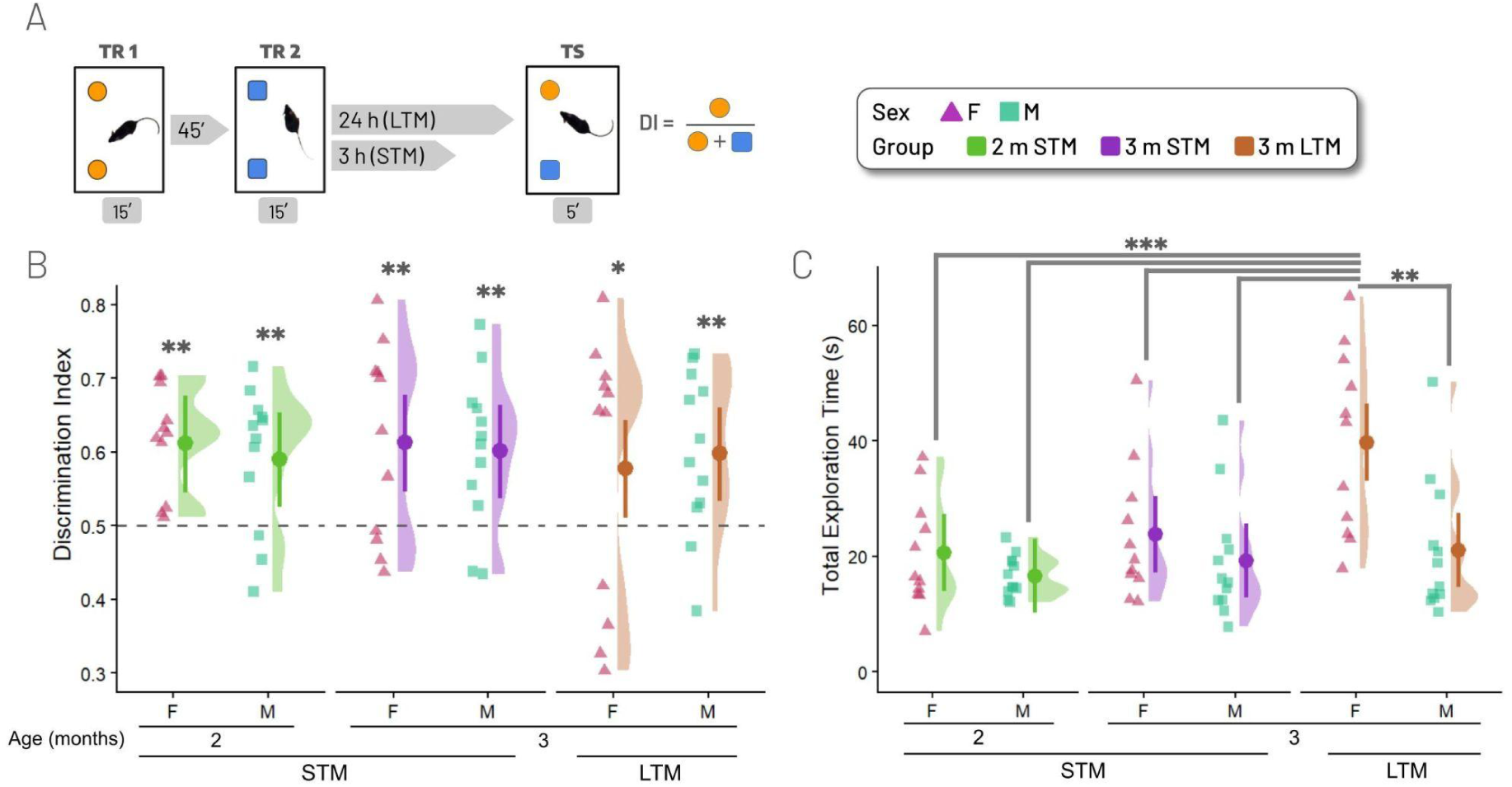
Characterization of Temporal Order Memory (TOM) in 2- and 3-month-old mice. **(A)** Schematic representation of the TOM protocol and Discrimination Index (DI) calculation, indicating training sessions (TR1 and TR2) and test session (TS) time points (STM: Short-Term, 3 h; LTM: Long-Term, 24 h). **(B-C)** Raincloud plots showing DI **(B)** and total exploration time **(C)** for groups tested at 2 months old, 3 h (2 m STM, green); 3 months old, 3 h (3 m STM, purple); and 3 months old, 24 h (3 m LTM, brown). The dashed line indicates the 0.5 chance level. Individual symbols represent single animals (blue squares for males, pink triangles for females). Crossbars indicate Estimated Marginal Means (EMMs) ± 95% CI derived from the GLMM. *, p < 0.05; **, p < 0.01; ***, p < 0.001 (Tukey’s post-hoc test).

Object exploration was quantified using the automated behavioral analysis pipeline RAINSTORM (D’hers et al., 2025). Artificial neural networks trained on manual annotations provided by five expert human scorers were used to identify exploratory behavior and automatically estimate object exploration times during TR1, TR2, and TS. Exploration times during TR1 and TR2 revealed no apparent differences between object/side (Figure S1), indicating the absence of intrinsic object bias.

A first cohort of mice (see Table 1) was used to evaluate TOM performance after short-term retention intervals at both 2 and 3 months of age, and after a long-term retention interval at 3 months of age. No significant effects of sex (χ^2^_(1)_ = 0.21, p = 0.6451), group (e.g., age and TS interval; χ^2^_(2)_ = 0.7, p = 0.7042), or sex-by-group interaction (χ^2^_(2)_ = 0.42, p = 0.8094) were detected in a type III ANOVA, indicating that both males and females performed similarly in this task at both ages and retention intervals. Moreover, Tukey’s post-hoc test confirmed that discrimination performance was significantly above chance level (DI = 0.5) in each of the three TS (Figure 1B, Table 3 in supplementary file), indicating that mice preferentially explored the older object in all conditions.

However, regarding total exploration time during TS (Figure 1C), a type III ANOVA revealed significant effects of the interaction between sex and group (χ^2^_(2)_ = 10.47, p = 0.0053). Tukey post-hoc comparisons revealed that 3-month-old females exhibited significantly higher total exploration times during LTM testing compared to all other sex/group pairs (Figure 1C).

Having established that 3-month-old mice exhibited robust TOM retention over a 24 h interval, we next evaluated long-term TOM performance in 2-month-old mice using an independent cohort (Table 1, Figure 2). No significant effects of sex (χ^2^_(1)_ = 0.18, p = 0.6703) were detected in a type III ANOVA, and discrimination index was significantly above chance level (DI = 0.5) in both males (13.6 % above chance, z = 3.83, p = 0.0001) and females (11.4 % above chance, z = 2.62, p = 0.0089; Figure 2A).

**Figure 2:**
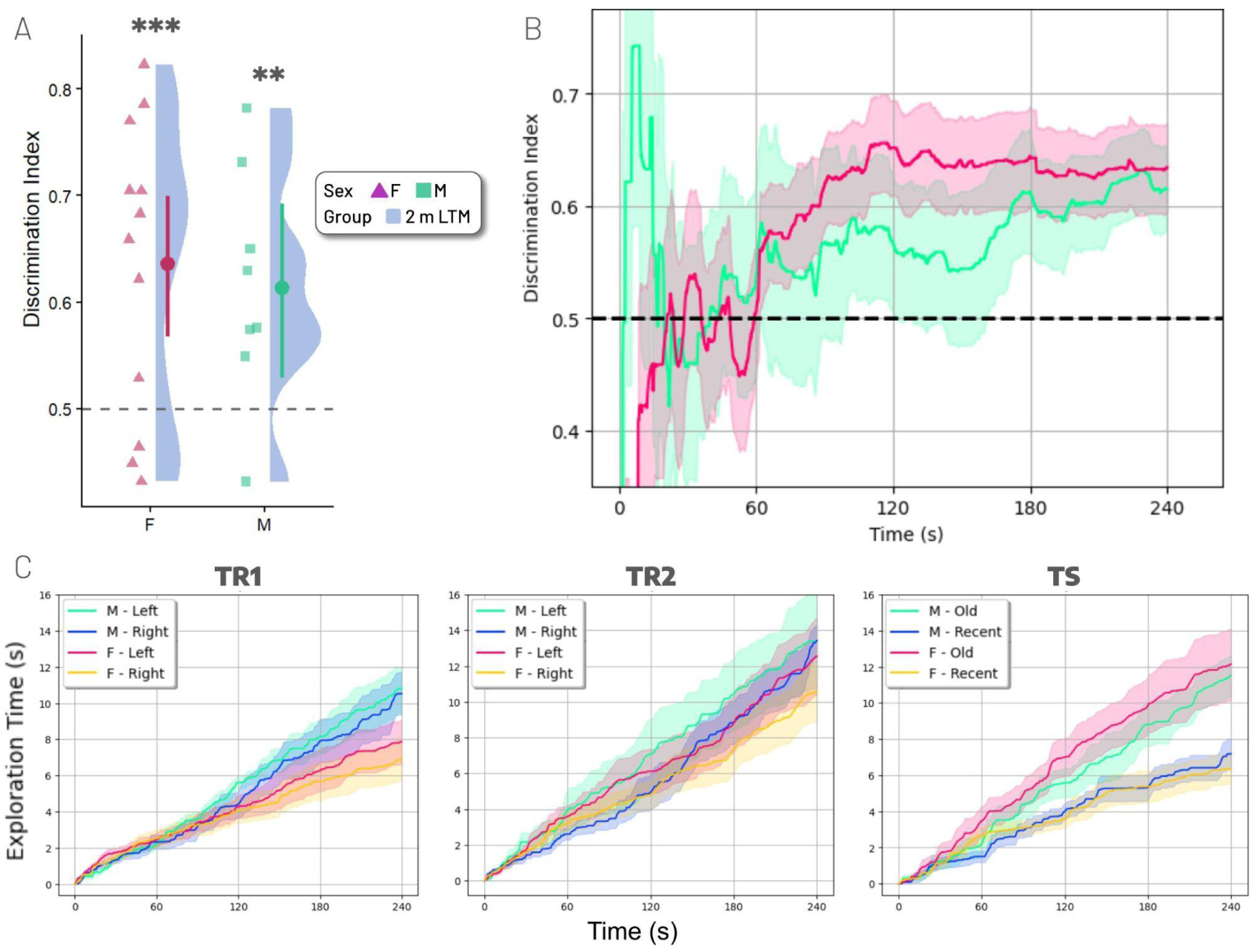
Characterization of long-term (24 h) Temporal Order Memory (TOM) in 2-month-old mice. **(A)** Raincloud plots showing the data distribution of the discrimination index (DI) for 2-month-old males and females tested at 24 h. The dashed line indicates the 0.5 chance level. Individual data symbols represent single animals (blue squares, males; pink triangles, females). Crossbars indicate Estimated Marginal Means (EMMs) ± 95% CI derived from the GLMM. **, p < 0.01; ***, p < 0.001 (Tukey’s post-hoc test). **(B)** Continuous dynamics of DI throughout TS obtained using RAINSTORM. Solid lines represent the mean DI over time, the shaded regions represent the SEM, and the dashed line represents the chance level (DI = 0.5). **(C)** Cumulative object exploration time throughout TR1 (left), TR2 (centre) and TS (right), separated by sex and either object location (left/right) during TR1 and TR2, or object identity (old/recent) during TS. Solid lines represent mean object exploration time and shaded regions indicate the SEM.

A detailed assessment of DI dynamics throughout the test session revealed a progressive emergence of temporal order discrimination in both sexes (Figure 2B). Consistent with this pattern, cumulative exploration curves showed a preferential exploration of the older object during testing, while no clear object/location preference was observed during training (Figure 2C). Comparable temporal dynamics were consistently observed across all TOM test sessions analyzed, irrespective of age or retention interval (Figure S1).

### 3.2. ERK2 phosphorylation dynamics during TOM

To investigate the molecular mechanisms underlying TOM, we characterized ERK1/2 phosphorylation dynamics in both the PFC and the HIP. Mice were either placed on the empty arena (HAB), trained once (TR1) or twice (TR2), and then sacrificed at 0, 1, or 2 h following the first session. Brain tissue was dissected and processed to obtain purified cytosolic- and nuclear-enriched protein fractions (Figure 3A). Analysis of pERK1/2 and total ERK1/2 immunoreactivity following SDS-PAGE (Figure 3B) revealed that, although the antibodies detected both ERK1 (44 kDa) and ERK2 (42 kDa), the ERK2 band exhibited the most consistent experience-dependent modulation. Therefore, subsequent analyses focused on ERK2 phosphorylation. No significant changes in ERK2 activation were detected in either the nuclear (Figure S2) or cytosolic (Figure S3) fractions of the HIP. Nuclear and cytosolic fractions from PFC evidenced similar activation patterns; since significant experience-dependent modulations of ERK2 phosphorylation were primarily identified in the cytosolic fraction of the PFC, the main focus was placed there.

**Figure 3:**
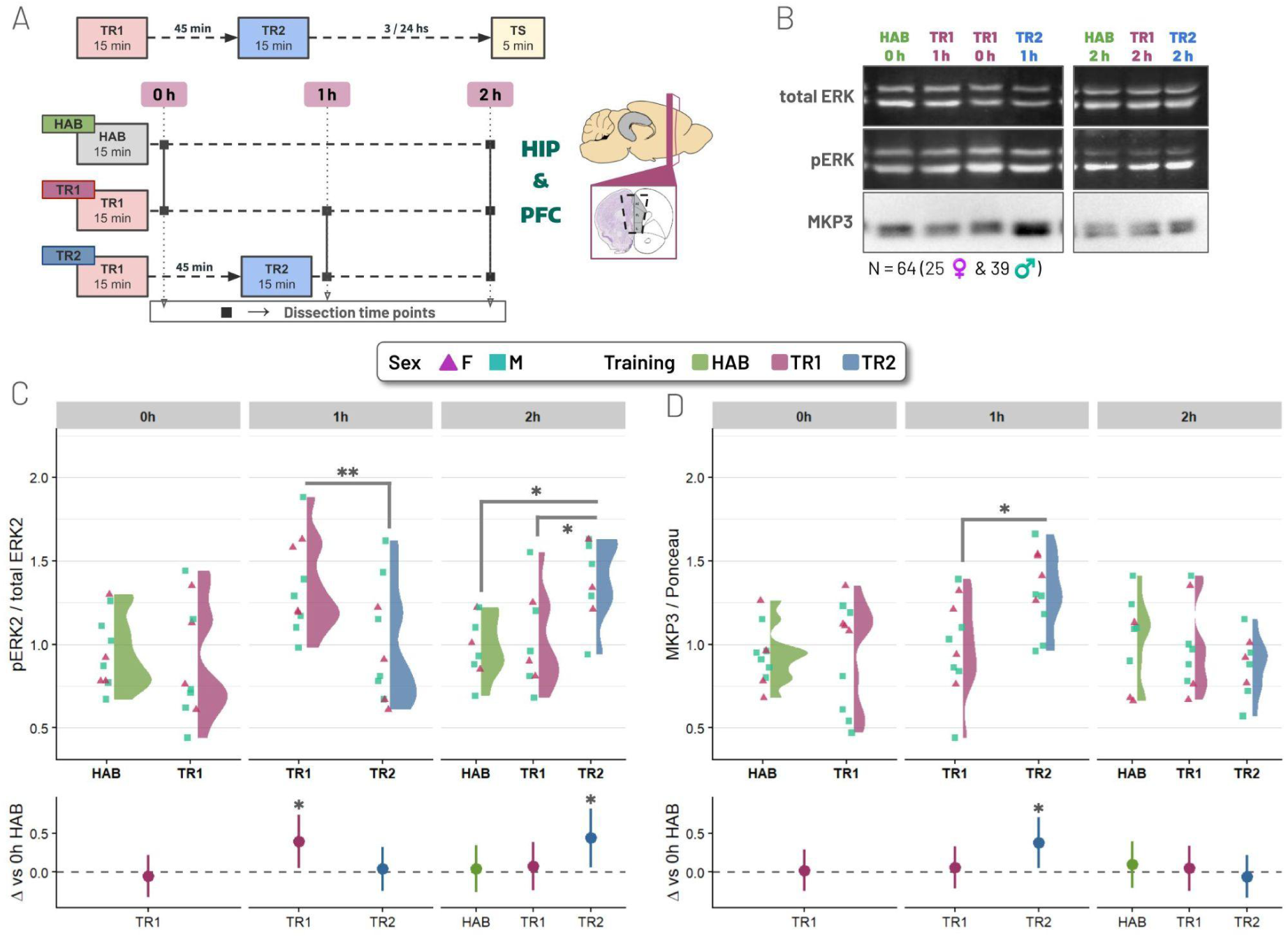
Experience-dependent cytosolic ERK2 activation and MKP3 expression in the prefrontal cortex during TOM acquisition. **(A)** Schematic representation of the training protocol and dissection time points (0, 1, and 2 h). **(B)** Representative immunoblots for phosphorylated ERK1/2 (pERK), total ERK1/2 (total ERK), and total MKP3 protein. Quantitative analysis was performed on the 42-kDa band (ERK2). **(C & D)** Raincloud plots depicting normalized ratio of cytosolic pERK2/total ERK2 **(C)** and MKP3 protein expression **(D)**. Colors represent: habituation to an empty arena (green, HAB), a single training session (purple, TR1), or two training sessions (blue, TR2). Individual data points represent individual subjects (blue squares, males; pink triangles, females). Crossbars indicate Estimated Marginal Means (EMMs) ± 95% CI derived from the GLMM. *, p < 0.05; **, p < 0.01 (Tukey’s post-hoc test).

Using a Gamma regression model, we found a significant interaction between training condition and time in a type II ANOVA (χ^2^_(2)_ = 15.15, p = 0.0005). Specifically, a single exposure to novel objects (TR1) induced a significant increase in ERK2 activation 1 hour after training (TR1_1h_vs HAB_0h_, z = 2.94, p = 0.0178), while no significant difference was found immediately after training (TR1_0h_ vs HAB_0h_, z = −0.51, p = 0.9636). Interestingly, when animals were subjected to the second training session (TR2), the initial phosphorylation event observed at 1 h was suppressed (TR2_1h_ vs HAB_0h_, z = 0.35, p = 0.9877; TR1_1h_ vs TR2_1h_, z = 2.61, p = 0.0091; Figure 3C). However, a delayed and significant increase in ERK2 activation emerged 1 h after TR2 relative to habituated controls at the same time point (TR2_2h_ vs HAB_2h_, z = 2.58, p = 0.0264; Figure 3C). We also assessed total ERK2 protein as a measure of ERK2 expression; however, no changes were found in any of the groups (Figure S2 and S3). These results suggest an experience-dependent regulation of cytosolic ERK activation in the PFC during sequential learning.

#### 3.2.1. Training-induced modulation of MKP3 expression in the PFC

Given that cytosolic ERK1/2 activation can be negatively regulated by the dual-specificity phosphatase MKP3 among others (Caunt & Keyse, 2013), we then characterized its cytosolic expression levels in the PFC (Figure 3B and D). A Type II ANOVA revealed a significant interaction between Training and Time (χ^2^_(2)_ = 6.91, p = 0.0316). Post-hoc comparisons against the global baseline (HAB_0h_) showed that MKP3 expression remained largely stable across groups, but significantly increased immediately after TR2 (TR2_1h_ vs HAB_0h_; z = 3.00, p = 0.0145). Furthermore, pairwise comparisons within the same post-training interval revealed higher MKP3 levels immediately after TR2 rather than 1 h after TR1 (TR2_1h_ vs TR1_1h_; z = 2.49, p = 0.0127; Figure 3D). These results suggest a potential link between ERK2 signaling and MKP3 function during TOM acquisition.

#### 3.2.2. Experience-dependent temporal coupling between ERK phosphorylation and MKP3 expression

Experience-dependent dynamics of ERK activation and MKP3 expression appear to be inversely related. Thus, increased ERK2 activation is associated with lower MKP3 protein levels, and vice versa. Consequently, we investigated whether they exhibit coordination at the individual level. To this end, we performed a Pearson correlation analysis segmented by experimental group in the cytosolic fraction of the PFC.

Immediately following a single training session (TR1_0h_), we observed a significant positive correlation between ERK2 activation and MKP3 expression (R = 0.689, p = 0.027; Figure 4A). This result suggests a parallel mobilization of both the kinase and the phosphatase in response to the novel stimulus. However, this relationship was drastically different one hour after a second training session (TR2_2h_), observing a significant negative correlation (R = −0.773, p = 0.025; Figure 4A). This finding indicates that, only after two training sessions, higher levels of MKP3 are associated with a decrease in ERK phosphorylation, suggesting a functional recruitment of a negative feedback loop induced by training repetition. In contrast, habituated control groups (HAB) showed no significant associations at any time point evaluated (p > 0.05).

**Figure 4:**
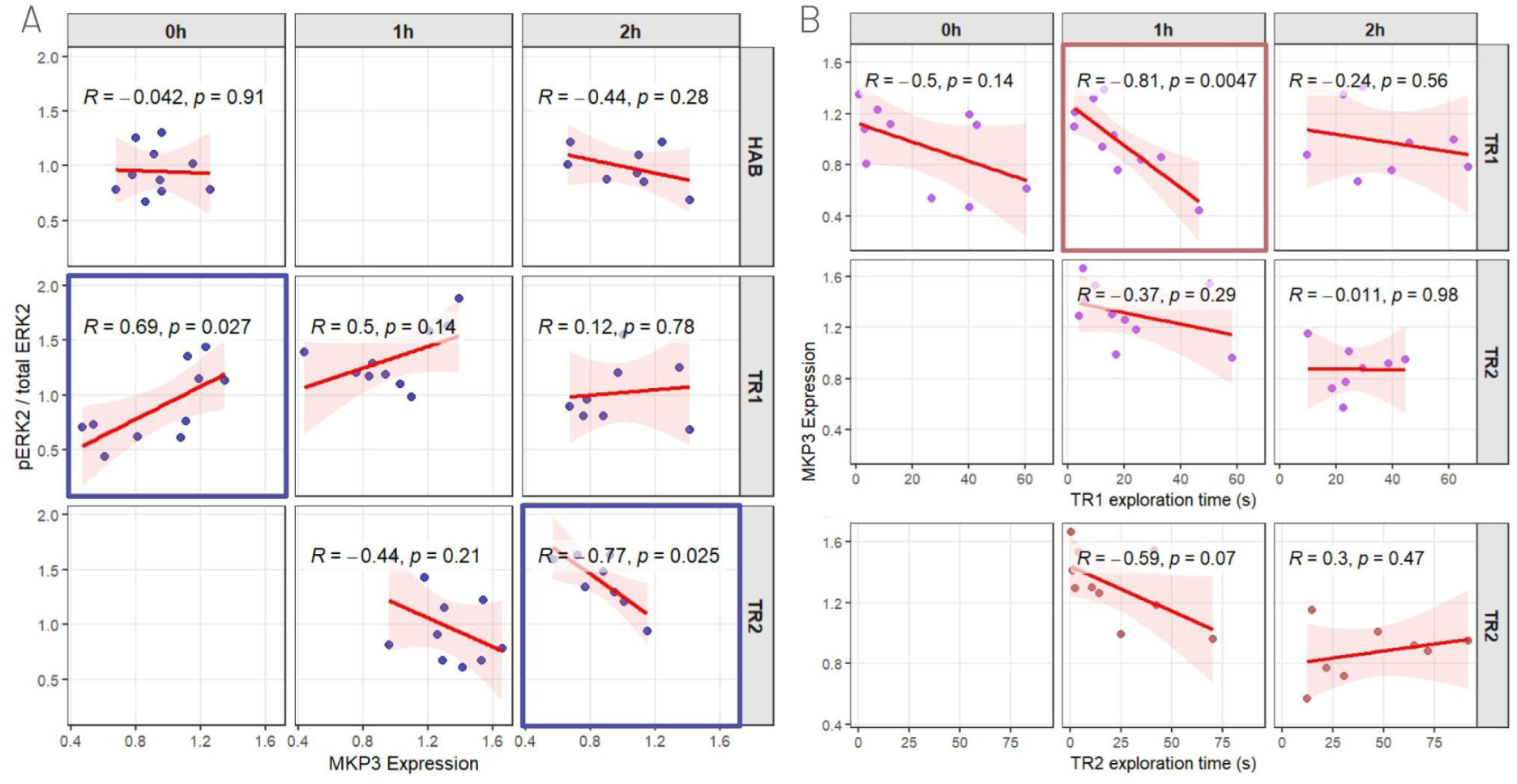
Correlation between MKP3 protein levels, ERK2 activation, and exploration time. **(A)** Pearson correlations between ERK2 activation and MKP3 expression in the PFC cytosolic fractions across conditions (training sessions and time). **(B)** Correlation between MKP3 expression and total exploration time. Each point represents an individual mouse. The red line represents the fitted linear regression, and the shaded area represents the 95% confidence interval. R: Pearson correlation coefficient. Blue & brown boxes highlight significant correlations (TR1_0h_ and TR2_2h_).

#### 3.2.3. Exploration time predicts MKP3 response magnitude but not ERK activation

To determine whether object exploration modulates the molecular response, we evaluated whether the total exploration time during TR1 and TR2 correlated with ERK2 activation and/or MKP3 protein levels. Interestingly, we found no significant correlations between exploration time and ERK2 activation at any time point (p > 0.05; Figure S4). In contrast, MKP3 expression did show a graded relationship with exploratory behavior. Specifically, in the TR1_1h_ group, a significant negative correlation was found between exploration time and MKP3 expression (R = −0.808, p = 0.0047; Figure 4B).

### 3.3. TeNOR reveals distinct memory traces

Finally, to delve into the TOM mechanism, we developed the Temporal Novel Object Recognition (TeNOR) protocol. This protocol mirrors the TOM training but diverges at the TS phase: mice were split into two groups, each tested 24 h later by exploring a completely novel object alongside either the object explored during TR1 (Prior_TR_) or during TR2 (Latter_TR_; Figure 5A).

**Figure 5:**
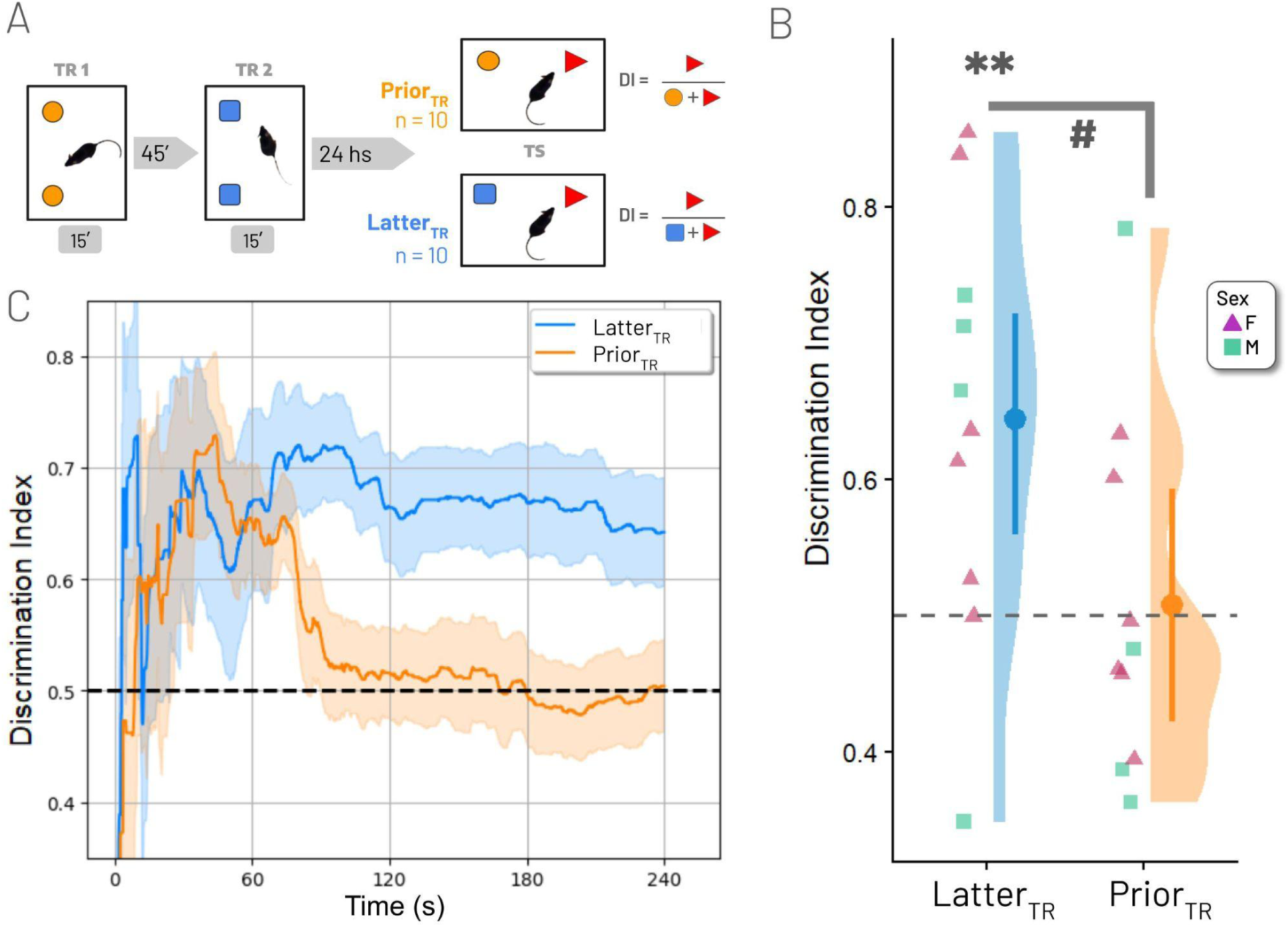
Differential recognition dynamics of old versus recent familiar objects in the TeNOR protocol. **(A)** Schematic representation of the TeNOR protocol. During TS, mice were presented with a novel object (red) alongside either the PriorTR object (orange) or the LatterTR object (blue). **(B)** Raincloud plot showing the global Discrimination Index (DI) for TS. The dashed line indicates the 0.5 chance level. Shapes map to sex (blue squares, males; pink triangles, females). Crossbars indicate Estimated Marginal Means (EMMs) ± 95% CI derived from the GLMM. #, p < 0.05; **, p < 0.01 (Tukey’s post-hoc test). **(C)** Continuous dynamics of the Discrimination Index throughout the testing session, analyzed via RAINSTORM. The solid lines represent the mean DI over time, the shaded regions represent the SEM, and the dashed line indicates the 0.5 chance level.

Our analysis revealed a significant main effect of the tested group (χ^2^ = 4.95, p = 0.0261; Figure 5B). Mice from Latter_TR_ robustly discriminated the familiar object against the novel one, exhibiting a strong preference for novelty (14.5 % above chance, z = 3.25, p < 0.0011). Mice from Prior_TR_ showed no significant discrimination against the novel object (0.8 % above chance, z = 0.18, p = 0.8609). Both groups exhibited similar exploration times (χ^2^ = 2.16, p = 0.1417).

Notably, continuous behavioral tracking using RAINSTORM enabled us to examine the temporal dynamics of the DI throughout the entire TS. This analysis revealed a brief initial period (approximately one minute) during which mice from Prior_TR_ preferentially explored the novel object over the familiar one, after which DI dropped to chance levels (Figure 5C). These findings suggest that memory of the old familiar object was preserved and capable of guiding behavior during the initial phase of testing. However, this effect was transient, suggesting that the relative influence of both memory representations on exploratory behavior changes rapidly over the course of testing.

## 4. Discussion

In this study, we characterized behavioral and molecular dynamics underlying Temporal Order Memory (TOM) in mice. Our findings support that TOM is a robust cognitive process that involves highly dynamic molecular signaling within the Prefrontal Cortex (PFC). Specifically, we identified a training-induced increase in ERK2 activation that is selectively modulated by subsequent experience and active, experience-dependent regulation mediated by the phosphatase MKP3. Furthermore, our results suggest that the preference for the older object in TOM may arise either from interference between competing memory traces or from differences in their relative stability during retrieval. The precise nature of TOM, whether it reflects the temporal discrimination of sequential events or the behavioral readout of memory interference, remains an open question.

Interestingly, a related form of retroactive interference has been described in an object-in-context task, where a second experience occurring one hour after the first disrupts long-term memory formation in a PFC-dependent manner (Martínez et al., 2014). The authors proposed that temporally proximal memories may compete for common cellular resources during consolidation, highlighting the PFC as a key structure in regulating interactions between memory traces. Furthermore, the idea that the PFC regulates competition between memory representations is consistent with previous evidence showing that this structure contributes to the control of interference during retrieval of recognition memories (Bekinschtein et al., 2013; Morici et al., 2015). In these studies, the PFC was proposed to bias behavioral output toward the most relevant memory trace among competing alternatives. Also, Autore and colleagues (2023) recently demonstrated that retroactive interference can promote adaptive forgetting through competition of newly formed engrams. In line with this framework, TOM performance may not necessarily reflect a timestamping mechanism. Instead, it may emerge from a competitive process in which consolidation of a first recognition memory is affected by a second, temporally close experience. Under this view, both object representations may be retained, but their behavioral expression is determined by the relative accessibility of the competing memory traces and by the mechanisms through which the PFC regulates their interaction. Whether this brain area actively allocates temporally proximal memories to competing neuronal or molecular ensembles, thereby shaping their subsequent behavioral expression, deserves further investigation.

### 4.1. ERK2 signaling in the Prefrontal Cortex

The Extracellular Signal-Regulated Kinase (ERK) pathway is a well-established requirement for long-term memory (LTM) across species (Atkins et al., 1998; Blum et al., 1999; Feld et al., 2005; Martin et al., 1997; Purcell et al., 2003; Ribeiro et al., 2005; Sweatt, 2004; among others), and object recognition memory stabilization has been specifically shown to require ERK activation (Kelly et al., 2003). Our results show that a single object exploration session (TR1) induces a significant increase in ERK2 phosphorylation in the PFC one hour later. Remarkably, a second training session (TR2) administered 45 min after TR1 effectively “reverses” or suppresses this peak at the 1-h time point, followed by a delayed increase in activation of ERK2 at the 2-h time point (Figure 6), indicating that ERK2 signaling is highly sensitive to exploratory experience and dynamically regulated by the interaction between temporally proximal memory-encoding events.

**Figure 6:**
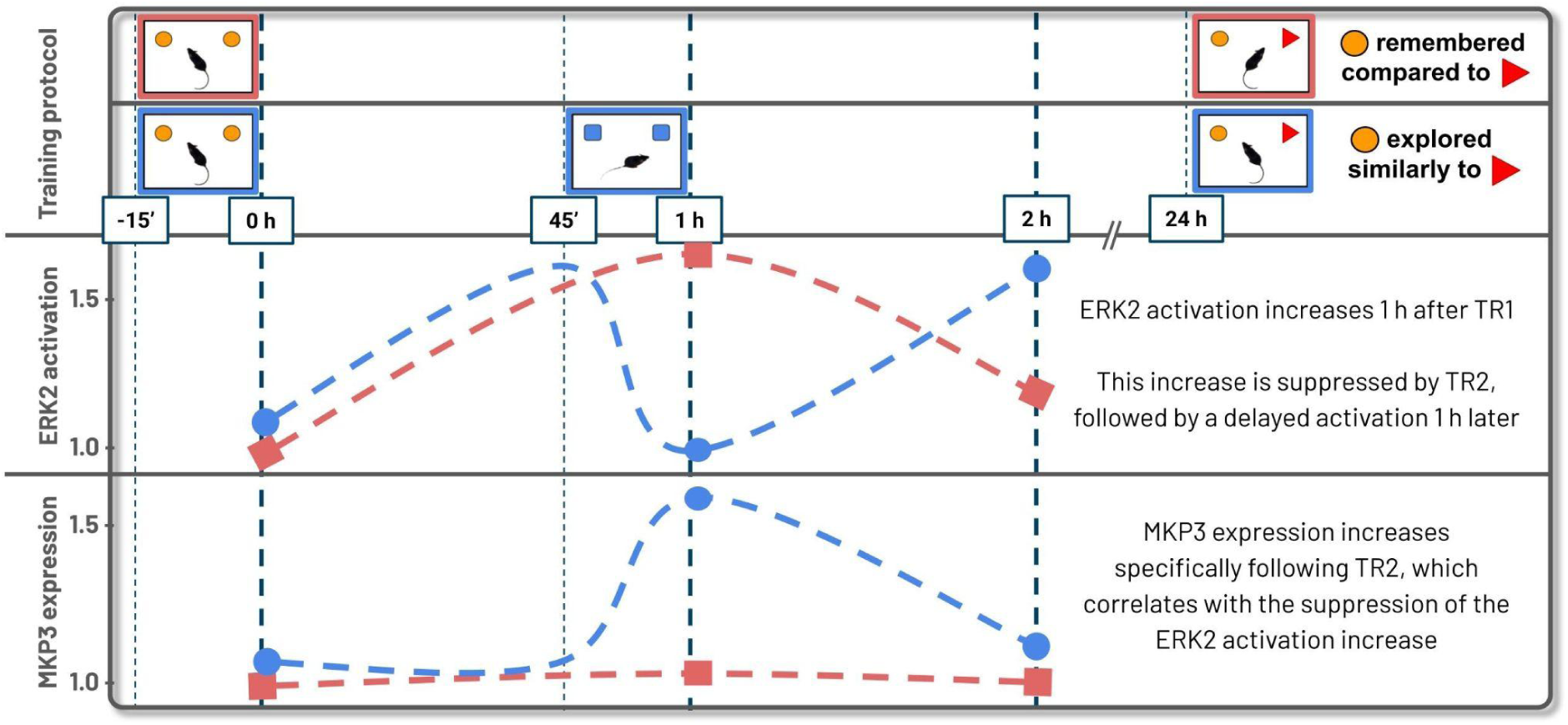
Working model summarizing the temporal dynamics of ERK2 phosphorylation and MKP3 expression following TR1 and TR2, highlighting an early transient ERK2 activation after TR1 and a delayed activation after TR2.

This rapid, experience-dependent reset of ERK2 activation is highly reminiscent of the findings by Cattaneo et al. (2020) in *Drosophila*, where a second learning trial erased the increase in ERK activation induced by the first one. By resetting ERK2 activity, the system may prevent a “molecular overlap” between separate events, thereby promoting the formation of distinct memory representations. Together with the behavioral evidence discussed above, these findings raise the possibility that dynamic ERK2 regulation contributes to the interaction and competition between coexisting memory traces. However, in the absence of information regarding the allocation of these signaling events within the brain, we cannot rule out the possibility that separate neural circuits compete for behavioral control.

Interestingly, in the absence of TR2, ERK2 activation eventually drops to baseline levels by the 2-h time point. This late decline does not appear to correlate with an upregulation of MKP3, suggesting that the spontaneous termination of ERK2 signaling after a single training experience may rely on constitutive dephosphorylation mechanisms or alternative phosphatase activity.

### 4.2. MKP3 expression in TOM

Feedback inhibition of ERK1/2 signaling can be mediated either by rapid posttranslational modifications of pathway components or by the slower induction of genes encoding pathway inhibitors, which requires de novo gene expression and protein synthesis (Lake et al., 2016). In support of this latter mechanism, Zhang and colleagues demonstrated that overexpression of wild-type MKP3 reduced EGF-induced ERK1/2 phosphorylation only after two hours of stimulation, whereas a phosphatase-dead mutant had no effect. A third possibility is that differential expression of scaffold proteins can also significantly modulate ERK1/2 dynamics. We observed a significant increase in MKP3 expression specifically following TR2 (TR2_1h_), which correlates with the suppression of ERK2 activation (Figure 3D). These findings support the possibility that MKP3 contributes either to delayed feedback regulation of ERK2 signaling or to experience-induced ERK2 suppression during TOM acquisition.

A particularly intriguing feature of this regulatory dynamic is the positive correlation between pERK2 and MKP3 immediately following the first training session (TR1_0h_, Figure 4A). While MKP3 is a canonical negative regulator of ERK—typically induced by ERK signaling to establish a negative feedback loop (Lake et al., 2016)—the immediacy of this correlation is unexpected. Because this parallel increase happens immediately after exploration, it is too fast to rely on *de novo* gene transcription. Instead, this early positive correlation likely reflects rapid translational control. The ERK pathway is known to regulate translation by phosphorylating 4E-BP1, promoting its dissociation from the translation initiation factor eIF4E (Herbert et al., 2002). By releasing this translational block, the initial surge in pERK2 can immediately drive the synthesis of MKP3 from pre-existing mRNA pools. Under this framework, the early positive correlation does not contradict MKP3’s role as a silencer, but rather captures the beginning of the feedback loop. The system mobilizes the kinase and its regulator in parallel as a preparatory step.

Finally, the transition to a strong negative correlation (TR2_2h_, Figure 4A) suggests that after repeated training, the phosphatase becomes effectively coupled to its target, ensuring the termination of the signal and laying the molecular groundwork required to compartmentalize temporally close experiences. Further supporting this regulated response is the negative correlation between exploration time during TR1 and MKP3 expression one hour later (TR1_1h_, Figure 4B). While it might seem counterintuitive that a longer exploration time is associated with lower levels of this phosphatase, this relationship could reflect a permissive window for memory consolidation. A prolonged sensory experience may require more sustained kinase activity to process the memory trace, potentially delaying the upregulation of MKP3. Conversely, brief exploratory events might trigger an early deployment of the phosphatase, truncating the signaling cascade earlier. Under this view, the MKP3 response could help scale the duration of the molecular trace to the intensity of the behavioral experience.

### 4.3. TeNOR: Insights on competing memories

To address whether the preference for the earlier-explored (Prior_TR_) object in TOM reflects a failure to recognize it, we developed the TeNOR (Temporal Novel Object Recognition) protocol. We found that while the memory of the recent object (Latter_TR_) drove sustained and robust novelty-seeking behavior throughout the test, the trace for the earlier-explored object appeared highly vulnerable. Interestingly, mice in the Prior_TR_ group exhibited a brief, initial period of discrimination (lasting approximately one minute) before their discrimination performance dropped to chance levels.

This initial discrimination suggests that the memory trace for the first object has not been completely erased; it remains available but possesses a diminished capacity to guide sustained behavior. This behavioral dynamic fits perfectly within the retroactive interference framework: TR2 interferes with the ongoing consolidation of the object memory from TR1, a process reflected at the molecular level by the MKP3-driven premature suppression of the TR1-induced ERK2 activation peak. In this context, the temporal preference seen in TOM could be an emergent property of two co-existing traces with different relative competitive strengths, rather than a strict binary state of remembering versus forgetting.

### 4.4. Limitations and Scope

While our results provide a clear molecular correlate for TOM, some limitations must be considered. First, our molecular assays were performed on whole-tissue extracts; while this identifies a global signaling state, it does not account for the potential heterogeneity of neuronal populations (e.g., pyramidal neurons vs. interneurons) that may play distinct roles in temporal processing. Furthermore, while the correlation between behavior, ERK2 activation, and MKP3 expression is strong, further pharmacological or genetic interventions—specifically targeting MKP3 or ERK2 in the PFC during the inter-trial interval—would be required to establish not only a definitive causal link between this molecular reset and the behavioral preference, but also the specific PFC circuitry involved in this type of memory.

## 5. Conclusion

In conclusion, we propose that Temporal Order Memory in mice relies on the Prefrontal Cortex’s ability to actively regulate object recognition memory interference to discriminate sequential events. The orchestrated activation of ERK2 and its rapid, experience-dependent downregulation—which correlates with increased MKP3 expression—provides a molecular framework for resolving competing memory traces. This mechanism, likely conserved from invertebrates to mammals, represents a fundamental computation through which the brain parses and organizes the continuous history of its experiences.

## Abbreviations

**DI** Discrimination Index

**ERK1/2** Extracellular Signal-Regulated Kinases 1 and 2

**GLMM** Generalized Linear Mixed Model

**HAB** Habituation

**HIP** Hippocampus

**ICC** Intraclass Correlation Coefficient

**LTM** Long-Term Memory

**MAPK** Mitogen-Activated Protein Kinase

**MKP3** Mitogen-Activated Protein Kinase Phosphatase 3

**PFC** Prefrontal Cortex

**RAINSTORM** Real and Artificial Intelligence for Neuroscience – Simple Tracker for Object Recognition Memory

**STM** Short-Term Memory

**TeNOR** Temporal Novel Object Recognition

**TOM** Temporal Order Memory

**TR** Training Session

**TS** Test Session.

## Author contributions

SD: Conceptualization, Methodology, Software, Validation, Formal Analysis, Investigation, Data Curation, Writing - Original Draft, Writing - Review & Editing, Visualization.

SOR and ADR: Methodology, Formal Analysis, Investigation, Writing - Review & Editing.

MF: Conceptualization, Methodology, Investigation, Supervision, Project Administration, Funding Acquisition, Resources, Writing - Original Draft, Writing - Review & Editing.

## Acknowledgments

The authors are grateful to Mariana Elias for her excellent assistance with mouse husbandry and animal care, and Maria Eugenia Cozzarin for her rigorous technical support.

## Declaration of generative AI and AI-assisted technologies in the writing process

During the preparation of this work, the authors used generative artificial intelligence (AI) tools, including large language models (LLMs), in order to improve language and readability of the manuscript. After using this tool/service, the first and corresponding authors reviewed and edited the content as needed, and they take full responsibility for the content of the publication.

## References

Adams, J. P., & Sweatt, J. D. (2002). Molecular Psychology: Roles for the ERK MAP Kinase Cascade in Memory. Annual Review of Pharmacology and Toxicology, 42(1), 135–163. 10.1146/annurev.pharmtox.42.082701.145401

Antoine, B., Serge, L., & Jocelyne, C. (2014). Comparative dynamics of MAPK/ERK signalling components and immediate early genes in the hippocampus and amygdala following contextual fear conditioning and retrieval. Brain Structure and Function, 219(1), 415–430. 10.1007/s00429-013-0505-y

Antunes, M., & Biala, G. (2012). The novel object recognition memory: Neurobiology, test procedure, and its modifications. Cognitive Processing, 13(2), 93–110. 10.1007/s10339-011-0430-z

Atkins, C. M., Selcher, J. C., Petraitis, J. J., Trzaskos, J. M., & Sweatt, J. D. (1998). The MAPK cascade is required for mammalian associative learning. Nature Neuroscience, 1(7), 602–609. 10.1038/2836

Autore, L., O’Leary, J. D., Ortega-de San Luis, C., & Ryan, T. J. (2023). Adaptive expression of engrams by retroactive interference. Cell Reports, 42(8), 112999. 10.1016/j.celrep.2023.112999

Barker, G. R. I., Bird, F., Alexander, V., & Warburton, E. C. (2007). Recognition Memory for Objects, Place, and Temporal Order: A Disconnection Analysis of the Role of the Medial Prefrontal Cortex and Perirhinal Cortex. The Journal of Neuroscience, 27(11), 2948–2957. 10.1523/JNEUROSCI.5289-06.2007

Barker, G. R. I., & Warburton, E. C. (2011). When Is the Hippocampus Involved in Recognition Memory? Journal of Neuroscience, 31(29), 10721–10731. 10.1523/JNEUROSCI.6413-10.2011

Bekinschtein, P., Renner, M. C., Gonzalez, M. C., & Weisstaub, N. (2013). Role of Medial Prefrontal Cortex Serotonin 2A Receptors in the Control of Retrieval of Recognition Memory in Rats. The Journal of Neuroscience, 33(40), 15716–15725. 10.1523/JNEUROSCI.2087-13.2013

Blum, S., Moore, A. N., Adams, F., & Dash, P. K. (1999). A Mitogen-Activated Protein Kinase Cascade in the CA1/CA2 Subfield of the Dorsal Hippocampus Is Essential for Long-Term Spatial Memory. The Journal of Neuroscience, 19(9), 3535–3544. 10.1523/JNEUROSCI.19-09-03535.1999

Brooks, M., E., Kristensen, K., Benthem, K., J. ,van, Magnusson, A., Berg, C., W., Nielsen, A., Skaug, H., J., Mächler, M., & Bolker, B., M. (2017). glmmTMB Balances Speed and Flexibility Among Packages for Zero-inflated Generalized Linear Mixed Modeling. The R Journal, 9(2), 378. 10.32614/RJ-2017-066

Cattaneo, V., San Martin, A., Lew, S. E., Gelb, B. D., & Pagani, M. R. (2020). Repeating or spacing learning sessions are strategies for memory improvement with shared molecular and neuronal components. Neurobiology of Learning and Memory, 172, 107233. 10.1016/j.nlm.2020.107233

Caunt, C. J., & Keyse, S. M. (2013). Dual-specificity MAP kinase phosphatases (MKPs): Shaping the outcome of MAP kinase signalling. The FEBS Journal, 280(2), 489–504. 10.1111/j.1742-4658.2012.08716.x

Datta, S. R., Anderson, D. J., Branson, K., Perona, P., & Leifer, A. (2019). Computational Neuroethology: A Call to Action. Neuron, 104(1), 11–24. 10.1016/j.neuron.2019.09.038

Dere, E., Dere, D., De Souza Silva, M. A., Huston, J. P., & Zlomuzica, A. (2018). Fellow travellers: Working memory and mental time travel in rodents. Behavioural Brain Research, 352, 2–7. 10.1016/j.bbr.2017.03.026

DeVito, L. M., & Eichenbaum, H. (2011). Memory for the Order of Events in Specific Sequences: Contributions of the Hippocampus and Medial Prefrontal Cortex. The Journal of Neuroscience, 31(9), 3169–3175. 10.1523/JNEUROSCI.4202-10.2011

D’hers, S., Robles, A. D., Ojea Ramos, S., Bollini, G. M., & Feld, M. (2025). RAINSTORM: Automated Behavioral Analysis for Mice Exploratory Behavior Using Artificial Neural Networks. Animal Behavior and Cognition. 10.1101/2025.04.07.647548

Eichenbaum, H. (2017). Memory: Organization and Control. Annual Review of Psychology, 68(1), 19–45. 10.1146/annurev-psych-010416-044131

Ennaceur, A., & Delacour, J. (1988). A new one-trial test for neurobiological studies of memory in rats. 1: Behavioral data. Behavioural Brain Research, 31(1), 47–59. 10.1016/0166-4328(88)90157-X

Farooq, A., & Zhou, M.-M. (2004). Structure and regulation of MAPK phosphatases. Cellular Signalling, 16(7), 769–779. 10.1016/j.cellsig.2003.12.008

Feld, M., Dimant, B., Delorenzi, A., Coso, O., & Romano, A. (2005). Phosphorylation of extra-nuclear ERK/MAPK is required for long-term memory consolidation in the crab. Behavioural Brain Research, 158(2), 251–261. 10.1016/j.bbr.2004.09.005

Feld, M., Galli, C., Piccini, A., & Romano, A. (2008). Effect on memory of acute administration of naturally secreted fibrils and synthetic amyloid-beta peptides in an invertebrate model. Neurobiology of Learning and Memory, 89(4), 407–418. 10.1016/j.nlm.2007.08.011

Hartig, F. (2026). DHARMa: Residual Diagnostics for Hierarchical (Multi-Level / Mixed) Regression Models. 10.32614/CRAN.package.DHARMa

Herbert, T. P., Tee, A. R., & Proud, C. G. (2002). The Extracellular Signal-regulated Kinase Pathway Regulates the Phosphorylation of 4E-BP1 at Multiple Sites. Journal of Biological Chemistry, 277(13), 11591–11596. 10.1074/jbc.M110367200

Istvan Lazar Jr., PhD; Istvan Lazar Sr., PhD, CSc. (n.d.). GelAnalyzer (Version 26.1) [Computer software]. Retrieved www.gelanalyzer.com

Janowsky, J. S., Shimamura, A. P., & Squire, L. R. (1989). Source memory impairment in patients with frontal lobe lesions. Neuropsychologia, 27(8), 1043–1056. 10.1016/0028-3932(89)90184-X

Kelly, Á., Laroche, S., & Davis, S. (2003). Activation of Mitogen-Activated Protein Kinase/Extracellular Signal-Regulated Kinase in Hippocampal Circuitry Is Required for Consolidation and Reconsolidation of Recognition Memory. The Journal of Neuroscience, 23(12), 5354–5360. 10.1523/JNEUROSCI.23-12-05354.2003

Krawczyk, M. C., Blake, M. G., Baratti, C. M., Romano, A., Boccia, M. M., & Feld, M. (2015). Memory reconsolidation of an inhibitory avoidance task in mice involves cytosolic ERK2 bidirectional modulation. Neuroscience, 294, 227–237. 10.1016/j.neuroscience.2015.03.019

Krawczyk, M. C., Navarro, N., Blake, M. G., Romano, A., Feld, M., & Boccia, M. M. (2016). Reconsolidation-induced memory persistence: Participation of late phase hippocampal ERK activation. Neurobiology of Learning and Memory, 133, 79–88. 10.1016/j.nlm.2016.06.013

Lake, D., Corrêa, S. A. L., & Müller, J. (2016). Negative feedback regulation of the ERK1/2 MAPK pathway. Cellular and Molecular Life Sciences, 73(23), 4397–4413. 10.1007/s00018-016-2297-8

Lenth, R. V., & Piaskowski, J. (2026). emmeans: Estimated Marginal Means, aka Least-Squares Means. https://rvlenth.github.io/emmeans/

Lopes, G., Bonacchi, N., FrazÃ£o, J., Neto, J. P., Atallah, B. V., Soares, S., Moreira, L., Matias, S., Itskov, P. M., Correia, P. A., Medina, R. E., Calcaterra, L., Dreosti, E., Paton, J. J., & Kampff, A. R. (2015). Bonsai: An event-based framework for processing and controlling data streams. Frontiers in Neuroinformatics, 9. 10.3389/fninf.2015.00007

Lüdecke, D., Ben-Shachar, M., Patil, I., Waggoner, P., & Makowski, D. (2021). performance: An R Package for Assessment, Comparison and Testing of Statistical Models. Journal of Open Source Software, 6(60), 3139. 10.21105/joss.03139

Martin, K. C., Michael, D., Rose, J. C., Barad, M., Casadio, A., Zhu, H., & Kandel, E. R. (1997). MAP Kinase Translocates into the Nucleus of the Presynaptic Cell and Is Required for Long-Term Facilitation in Aplysia. Neuron, 18(6), 899–912. 10.1016/S0896-6273(00)80330-X

Martínez, M. C., Villar, M. E., Ballarini, F., & Viola, H. (2014). Retroactive interference of object-in-context long-term memory: Role of dorsal hippocampus and medial prefrontal cortex. Hippocampus, 24(12), 1482–1492. 10.1002/hipo.22328

Mathis, A., Mamidanna, P., Cury, K. M., Abe, T., Murthy, V. N., Mathis, M. W., & Bethge, M. (2018). DeepLabCut: Markerless pose estimation of user-defined body parts with deep learning. Nature Neuroscience, 21(9), 1281–1289. 10.1038/s41593-018-0209-y

Mitchell, J. (1998). The medial frontal cortex and temporal memory: Tests using spontaneous exploratory behaviour in the rat. Behavioural Brain Research, 97(1–2), 107–113. 10.1016/S0166-4328(98)00032-1

Morici, J. F., Bekinschtein, P., & Weisstaub, N. V. (2015). Medial prefrontal cortex role in recognition memory in rodents. Behavioural Brain Research, 292, 241–251. 10.1016/j.bbr.2015.06.030

Nath, T., Mathis, A., Chen, A. C., Patel, A., Bethge, M., & Mathis, M. W. (2019). Using DeepLabCut for 3D markerless pose estimation across species and behaviors. Nature Protocols, 14(7), 2152–2176. 10.1038/s41596-019-0176-0

Ojea Ramos, S., Andina, M., Romano, A., & Feld, M. (2021). Two spaced training trials induce associative ERK-dependent long term memory in Neohelice granulata. Behavioural Brain Research, 403, 113132. 10.1016/j.bbr.2021.113132

Ojea Ramos, S., Medina, C., Krawczyk, M. D. C., Millan, J., Acutain, M. F., Romano, A. G., Baez, M. V., Urbano, F. J., Boccia, M. M., & Feld, M. (2026). Hippocampal Erk2 Dimerization is Critical for Memory Reconsolidation and Synaptic Plasticity&nbsp; SSRN. 10.2139/ssrn.6169711

Posit team. (2026). RStudio: Integrated Development Environment for R. Posit Software, PBC. http://www.posit.co/

Purcell, A. L., Sharma, S. K., Bagnall, M. W., Sutton, M. A., & Carew, T. J. (2003). Activation of a Tyrosine Kinase-MAPK Cascade Enhances the Induction of Long-Term Synaptic Facilitation and Long-Term Memory in Aplysia. Neuron, 37(3), 473–484. 10.1016/S0896-6273(03)00030-8

R Core Team. (2026). R: A Language and Environment for Statistical Computing. R Foundation for Statistical Computing. 10.32614/R.manuals

Ribeiro, M. J., Schofield, M. G., Kemenes, I., O’Shea, M., Kemenes, G., & Benjamin, P. R. (2005). Activation of MAPK is necessary for long-term memory consolidation following food-reward conditioning. Learning & Memory, 12(5), 538–545. 10.1101/lm.8305

Schneider, C. A., Rasband, W. S., & Eliceiri, K. W. (2012). NIH Image to ImageJ: 25 years of image analysis. Nature Methods, 9(7), 671–675. 10.1038/nmeth.2089

Sweatt, J. D. (2004). Mitogen-activated protein kinases in synaptic plasticity and memory. Current Opinion in Neurobiology, 14(3), 311–317. 10.1016/j.conb.2004.04.001

Thomas, G. M., & Huganir, R. L. (2004). MAPK cascade signalling and synaptic plasticity. Nature Reviews Neuroscience, 5(3), 173–183. 10.1038/nrn1346

Tulving, E. (2002). Episodic Memory: From Mind to Brain. Annual Review of Psychology, 53(1), 1–25. 10.1146/annurev.psych.53.100901.135114

