## Supplemental figures and Table 3 for "The pERKs of Temporal Order Memory in mice"

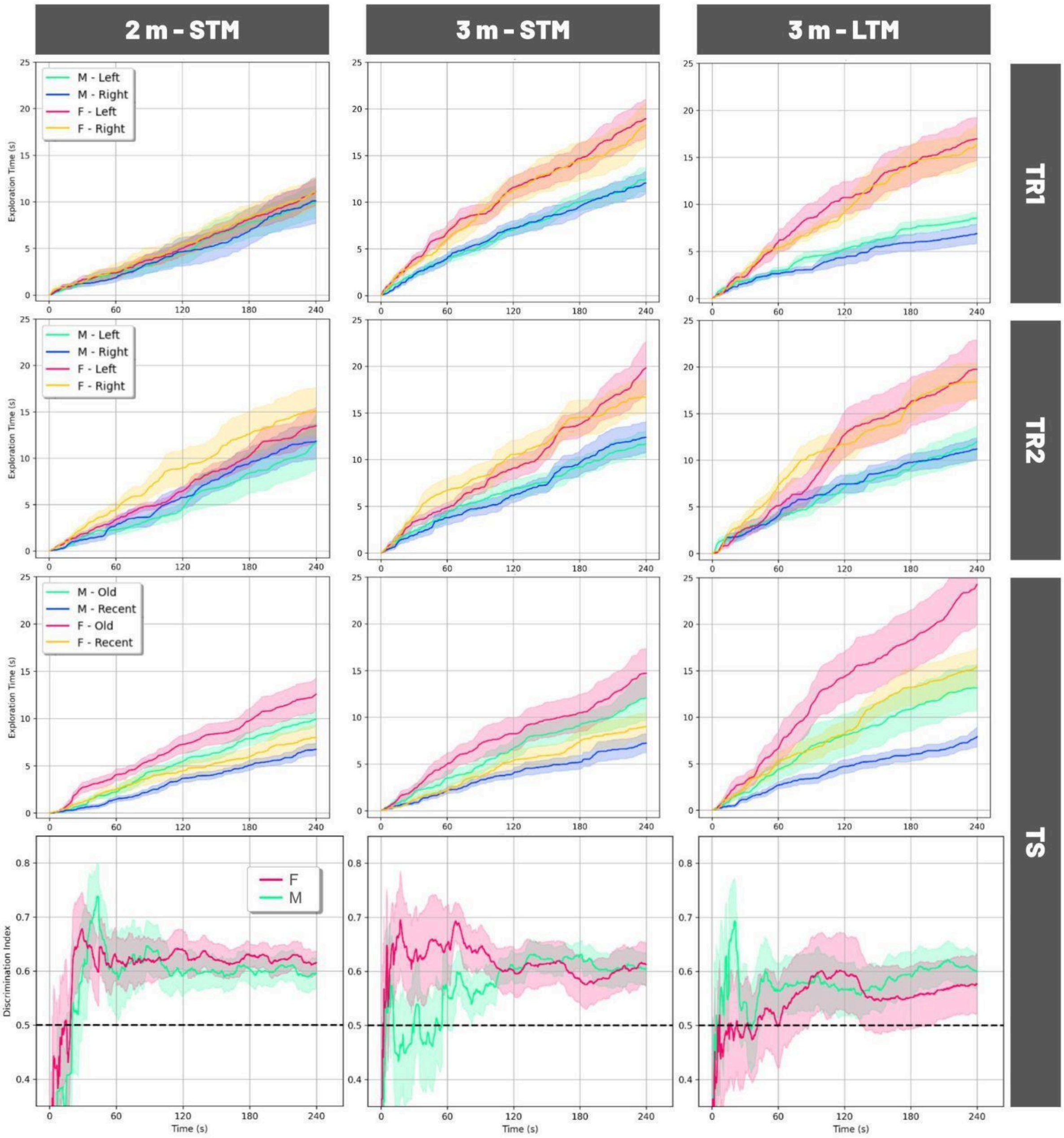


**Figure S1**: Exploration dynamics during TR1, TR2 and TS; and absence of intrinsic object bias during TR1 and TR2 in each experiment.

Cumulative object exploration time throughout TR1 (first row), TR2 (second row) and TS (third row), separated by sex and side of the arena for object location (left/right) during TR1 and TR2, or object identity (old/recent) during TS. Each experiment is shown in a different column. Solid lines represent mean object exploration time and shaded regions indicate the SEM.

Continuous dynamics of DI throughout TS obtained using RAINSTORM is shown in the last row. Solid lines represent the mean DI over time, the shaded regions represent the SEM, and the dashed line represents the chance level (DI = 0.5).

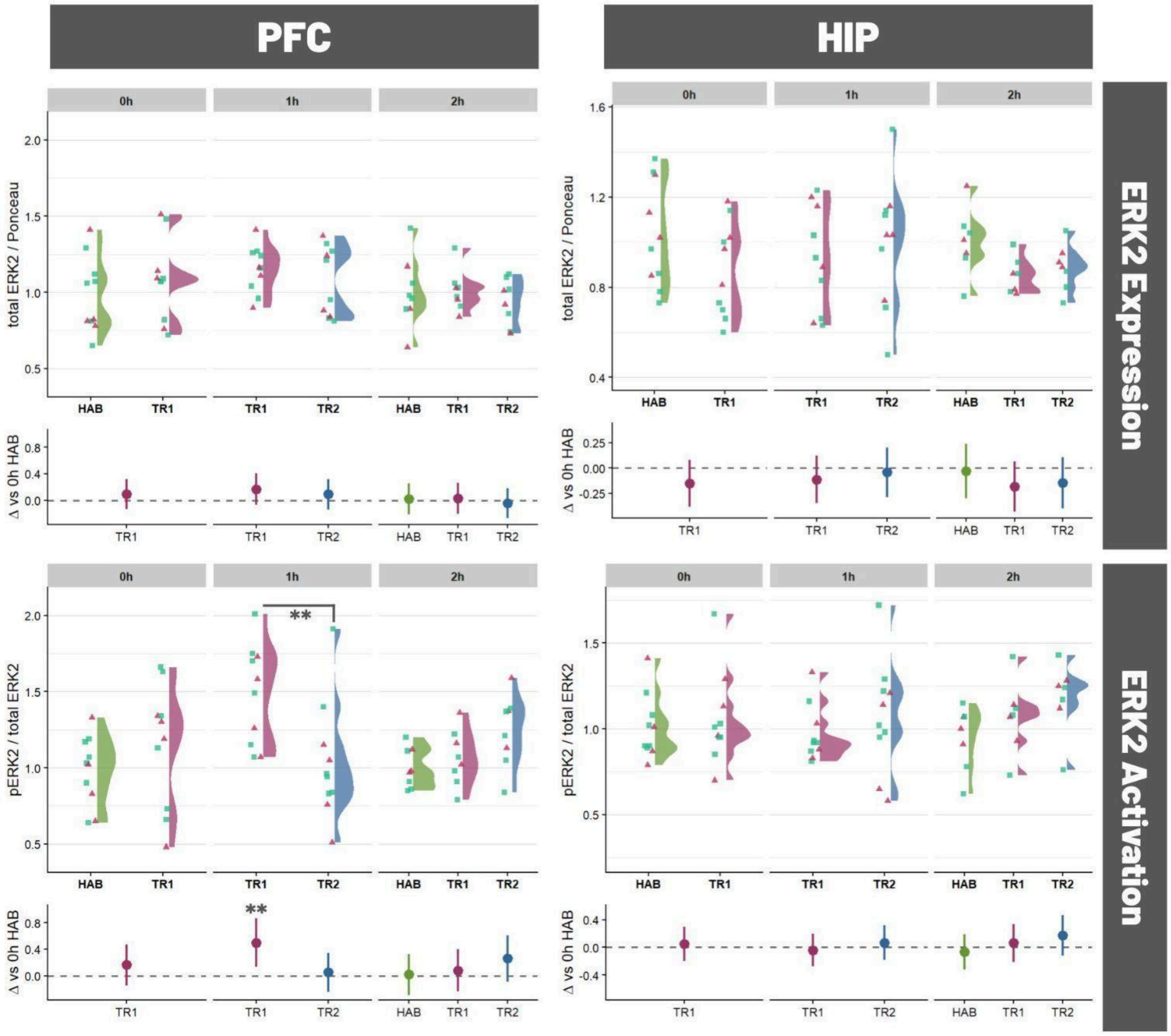


**Figure S2**: ERK2 activation and expression in nuclear fractions of PFC and HIP.
Colors represent: habituation to an empty arena (green, HAB), a single training session (purple, TR1), or two training sessions (blue, TR2). Individual data points represent individual subjects (blue squares, males; pink triangles, females). Crossbars indicate Estimated Marginal Means (EMMs) ± 95% CI derived from the GLMM. **, $p < 0.01$ (Tukey's post-hoc test).

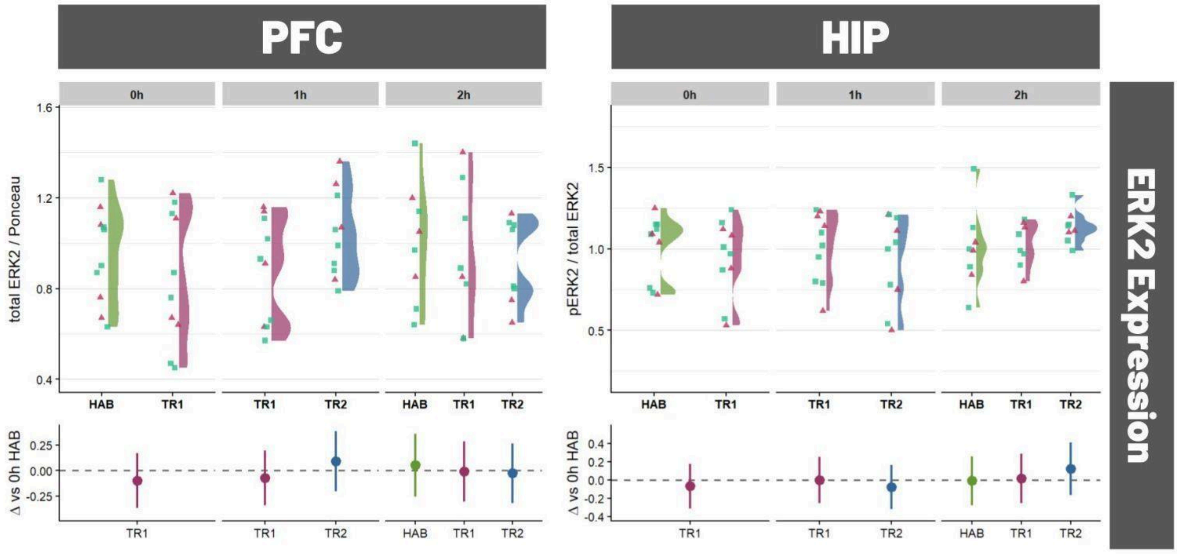


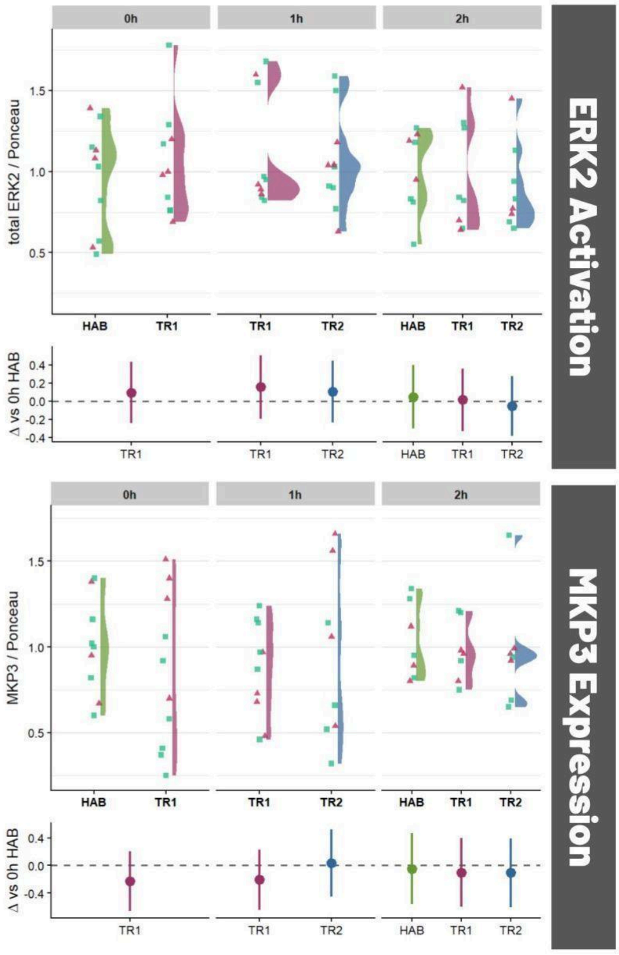


**Figure S3**: ERK2 activation, ERK2 expression, and MKP3 expression in cytosolic fractions.

Colors represent: habituation to an empty arena (green, HAB), a single training session (purple, TR1), or two training sessions (blue, TR2). Individual data points represent individual subjects (blue squares, males; pink triangles, females). Crossbars indicate Estimated Marginal Means (EMMs) ± 95% CI derived from the GLMM.

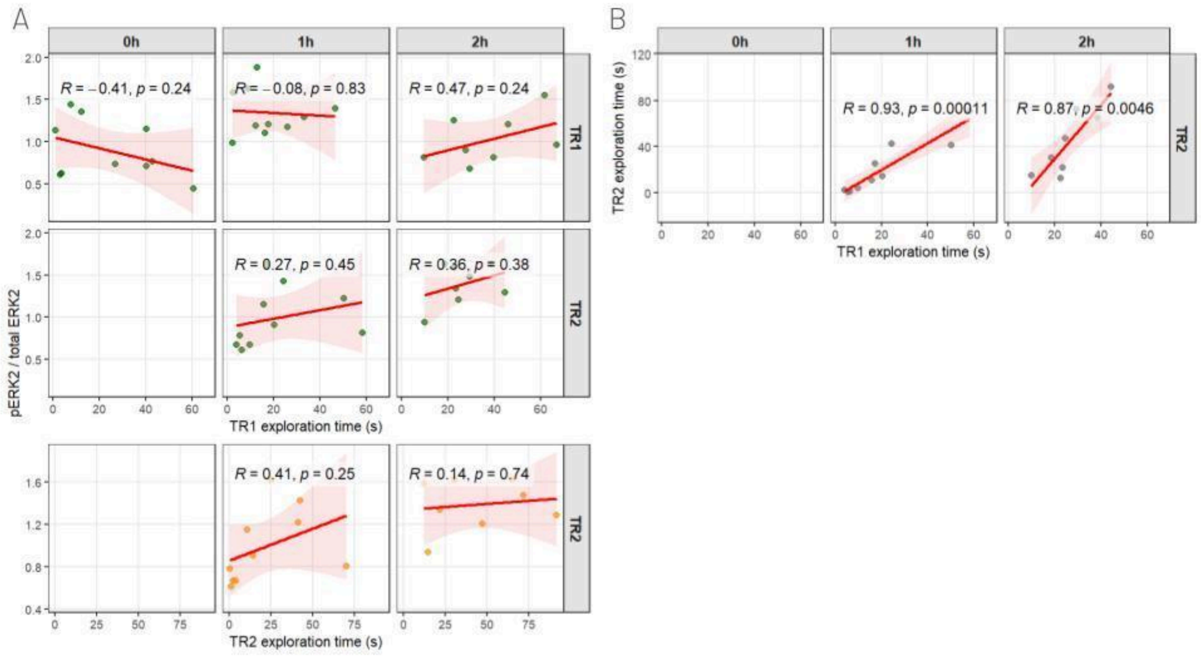


**Figure S4**: Correlation between ERK2 activation and total exploration time.
**(A)** Pearson correlations between ERK2 activation in the PFC cytosolic fraction and total exploration time across conditions (training sessions and time).
**(B)** Pearson correlations between total exploration time between TR1 and TR2.
Each point represents an individual mouse. The red line represents the fitted linear regression, and the shaded area represents the 95% confidence interval. R: Pearson correlation coefficient.

| group & sex | Δ DI (vs. Chance) | CI | z ratio | p value |
|---|---|---|---|---|
| 2 m STM - F | +11.3% | [4.5 ; 17.6] | 3.22 | 0.0013 |
| 3 m STM - F | +11.4% | [4.6 ; 17.8] | 3.26 | 0.0011 |
| 3 m LTM - F | +7.8% | [1 ; 14.4] | 2.26 | 0.0241 |
| 2 m STM - M | +9.1% | [2.6 ; 15.3] | 2.73 | 0.0064 |
| 3 m STM - M | +10.2% | [3.7 ; 16.3] | 3.05 | 0.0023 |
| 3 m LTM - M | +9.8% | [3.3 ; 16] | 2.94 | 0.0033 |

**Table 3**: Discrimination performance compared against chance level (DI = 0.5) for 2-month-old (2 m) and 3-month-old (3 m) female (F) and male (M) mice evaluated 3 h (STM) or 24 h (LTM) after TR2.
